# Equivalent fitness increase achieved by active learning-navigated habitat reconstruction and evolution-induced genome mutation

**DOI:** 10.64898/2026.03.29.715165

**Authors:** Zipeng Lu, Bei-Wen Ying

## Abstract

Fitness increase, impacted by genetic and environmental traits, requires evolutionary changes in genomes or ecological changes in habitats. Whether and how habitat reconstruction can compensate for the genetic changes remains unclear. The present study offers an experimental comparison of bacterial fitness increase *via* evolutionary and ecological strategies to verify that habitat reconstruction has the potential to avert genetic restriction, challenging the genetic-only view of adaptation. Six finely tuned media with different combinations and five evolved lineages with different mutations, both of which allowed equivalent increases in bacterial growth rates, were achieved through active machine learning and experimental evolution, respectively. Transcriptome changes accompanied by fitness increase were limited but divergent among the six fitted media, compared to changes that were broad but similar among the five fitted genomes. The universal transcriptome reorganization in fitted genomes and media was likely driven by surrounding metabolisms, indicating escape routes for genetic changes that increase fitness through habitat reconstruction.

## Introduction

Bacterial fitness is the output of the equations formed by genetic and environmental terms, such as the genome and the habitat. Nevertheless, it was extensively explored by genetics in adapting to a defined environment from an evolutionary viewpoint. The fitness landscape is employed to understand the relationship between genotype and fitness ^1,2^, facilitating insights into adaptation across various biological levels ^3^. Traditionally, adaptation is understood as a genetic process in which natural selection ^4^ favors beneficial mutations ^5,6^, resulting in fitness gain over time ^7,8^. This gene-centric view has dominated evolutionary biology for decades ^9,10^, extensively supported by the experimental evolution that demonstrated growth fitness increase driven by genetic changes under constant selection pressures ^11,12^.

However, environmental variance has been shown to shape evolution for fitness increase ^13,14^. Introducing environmental traits to the genetic changes frequently resulted in a canceling effect on growth fitness ^15,16^ and influenced the epistatic fitness landscape ^17,18^. From an ecological perspective, the benefits of improving fitness through habitat reconstruction were considered. Theoretical studies have demonstrated the significance of non-genetic traits, e.g., niche construction ^19,20^, where the environment could play a pivotal role in shaping adaptive trajectories for fitness increase ^19^. Adaptive niche construction has been shown to alter the distribution of environments and increase population fitness ^21^. It indicated that living organisms actively modify their environments (habitats) in ways that affect their fitness. However, experimental demonstrations of fitness increase purely through environmental optimization, without accompanying genetic change, have been scarce and limited by technical constraints in designing high-dimensional habitats. The role of environmental change in fitness changes has proven more challenging to identify and formalize ^22–24^.

Lately, advances in machine learning (ML) ^25–27^ offer a powerful solution to this challenge as a novel experimental avenue to explore how environmental reconstruction alone can drive fitness increase. ML-assisted habitat reconstruction enables fitness improvements achieved by optimizing environmental inputs (e.g., nutrient composition) without invoking genetic mutations. ML algorithms can navigate high-dimensional optimization problems, making them well-suited for designing complex systems ^28,29^. Employing ML in environmental reconstruction could not only adjust habitat parameters (e.g., nutrient composition) based on measured fitness outputs (e.g., growth rate) but also identify the decision-making elements for fitness increase. The decision-making factors for bacterial growth increase were addressed under well-controlled laboratory conditions (media) by decision tree machine learning algorithms ^16,30^. The ML-assisted approach identifies configurations that enhance fitness with greater efficiency than random or manually guided methods. It facilitates a new paradigm: the environment is iteratively optimized to fit the organism rather than evolving organisms to fit a fixed environment.

Whether the genetic changes for fitness increase by evolution can be averted or compensated for by habitat reconstruction? The present study sought to provide an experimental demonstration of whether evolutionary constraints could be overcome by habitat reconstruction, which is restricted to the balancing of the resources in the environment but not the synthetic reconstruction of the microbial community. Experimental evolution and ecology were employed to achieve the fitted genomes that conquered the genetic constraint and fitted habitats that averted the genetic constraint for fitness increase. Experimental evolution could be conducted through serial transfer, that is, repeated dilution of growing bacterial populations to mimic genome evolution, adapting to the defined environment. Our previous studies demonstrated that reduced growth fitness due to genome reduction in *Escherichia coli* can be compensated through experimental evolution ^31^, which provided the evolved genomes for the present study. Experimental ecology could be achieved by combining the high-throughput growth assay ^32^ for bacterial population dynamics ^33^ and active learning associated with machine learning ^34–36^. The active machine learning was employed here to mimic environmental discovery for fitted habitats.

## Results

### Experimental design of bacterial fitness increases via evolutionary and ecological strategies

An experimental demonstration of two fundamental strategies for bacterial fitness increase, experimental evolution to find fitted genomes and experimental ecology to find fitted habitats (Fig. 1A, magenta) were conducted. The original condition (Ori) was designated as a genome-reduced *E. coli* strain (Anc, see Materials and Methods) ^37^ growing in a chemically defined medium (M63) (Fig. 1A). Varying the chemical composition of M63 could change the growth rates of the *E. coli* strains carrying varied genome sizes ^38^. The evolutionary strategy for the fitness increase of Ori was to evolve Anc to obtain mutated genomes (EVOs) of increased growth rates in M63. The ecological strategy was to fine-tune M63 to obtain optimized media (OPTs) for increased growth rates of Anc.

**Figure 1.**
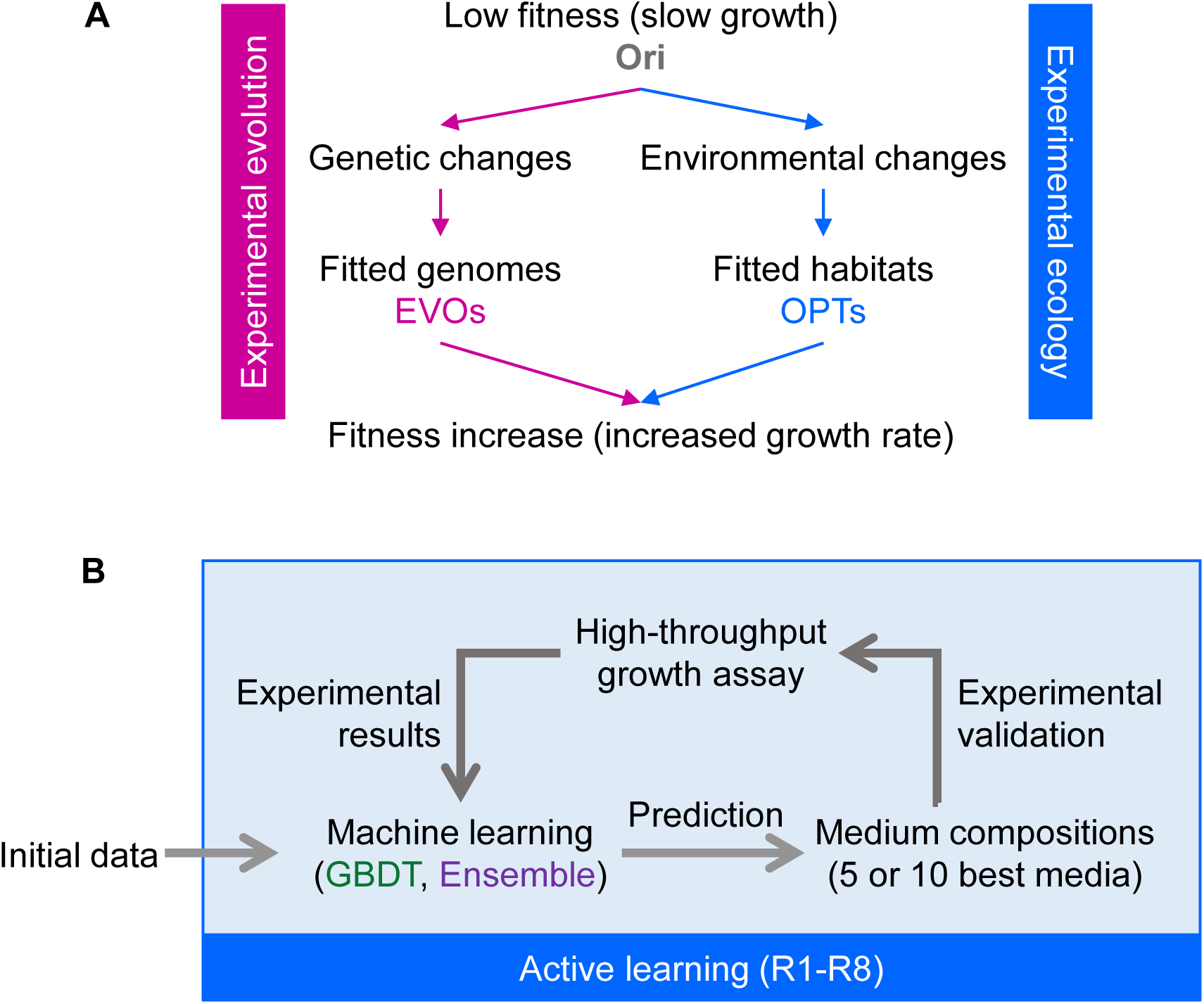
Scheme of evolutionary and ecological strategies for increased fitness. **A.** Experimental overview. Ori, EVOs, and OPTs represent the original genotype grown under the initial environment, the evolved genotypes remaining at the initial environment, and the original genotype grown under changed environments, respectively. Magenta and blue indicate evolutionary and ecological strategies, mimicked by experimental evolution and experimental ecology, respectively. **B.** Active learning for media optimization as an ecological strategy for habitat reconstruction. The main steps are outlined, i.e., ML model construction, medium prediction for increased growth, experimental validation by growth assay, and data integration to improve ML models.

Experimental evolution has been performed in our previous study ^31^. Anc was evolved in M63, and multiple evolved lineages (EVOs) were obtained (Fig. S1). Five out of nine EVOs, presenting the most significant fitness increase, were selected as fitted genomes for use in the present study. Experimental ecology was employed to achieve fitted habitats (OPTs) (Fig. 1A, blue) through the machine learning (ML)-guided fine-tuning of medium compositions in the present study (Fig. 1B). The medium (M63) used in the experimental evolution served as the initial habitat, comprising seven pure chemical compounds (Table S1). Fine-tuning the concentrations of these compounds to find fitted habitats (OPTs) enabled Anc to have increased growth rates equivalent to those of EVOs.

### ML-guided medium fine-tuning succeeded in finding fitted habitats

ML-guided fine-tuning of medium compositions was performed using two different models (Fig. 1B). One was the gradient-boosting decision tree (GBDT) model, as we previously used for medium optimization for bacterial ^35,39^ and mammalian cells^34,36^. The other was the Ensemble model, a combination of four different algorithms: GBDT, support vector regressor (SVR), k-nearest neighbors (k-NN), and neural networks (NN). To obtain the training data (Initial) for both ML models, the growth assay was initially performed under 99 different medium compositions. To enhance the efficiency of medium fine-tuning, active learning was employed, i.e., repeated model construction, prediction, and experimental verification steps (Fig. 1B). The active learning was conducted over eight rounds (R1-R8) using two ML models separately. Finally, a total of 1,350 growth assays in 209 different media were conducted through active learning.

Both ML models succeeded in gradually improving Anc’s growth as rounds proceeded, but presented distinct efficiencies. The growth rates under the Ensemble-predicted media increased gradually as the active learning rounds progressed and ultimately saturated at R5 (Fig. 2A). This trend of fitness increase and saturation was consistent across evaluations using the mean of biological replicates and individual assays (Fig. S2A). In comparison, the growth rates under the GBDT-predicted media presented both increases and decreases as rounds proceeded, which likely reached saturation at R3 (Fig. 2B, Fig. S2B). It revealed that the Ensemble model outperformed the GBDT model in fine-tuning the medium compositions for improved bacterial growth, as verified by their prediction accuracy, i.e., Ensemble was significantly better than GBDT (Fig. S3). The results demonstrated that active learning successfully achieved multiple habitats for Anc to have improved fitness.

**Figure 2.**
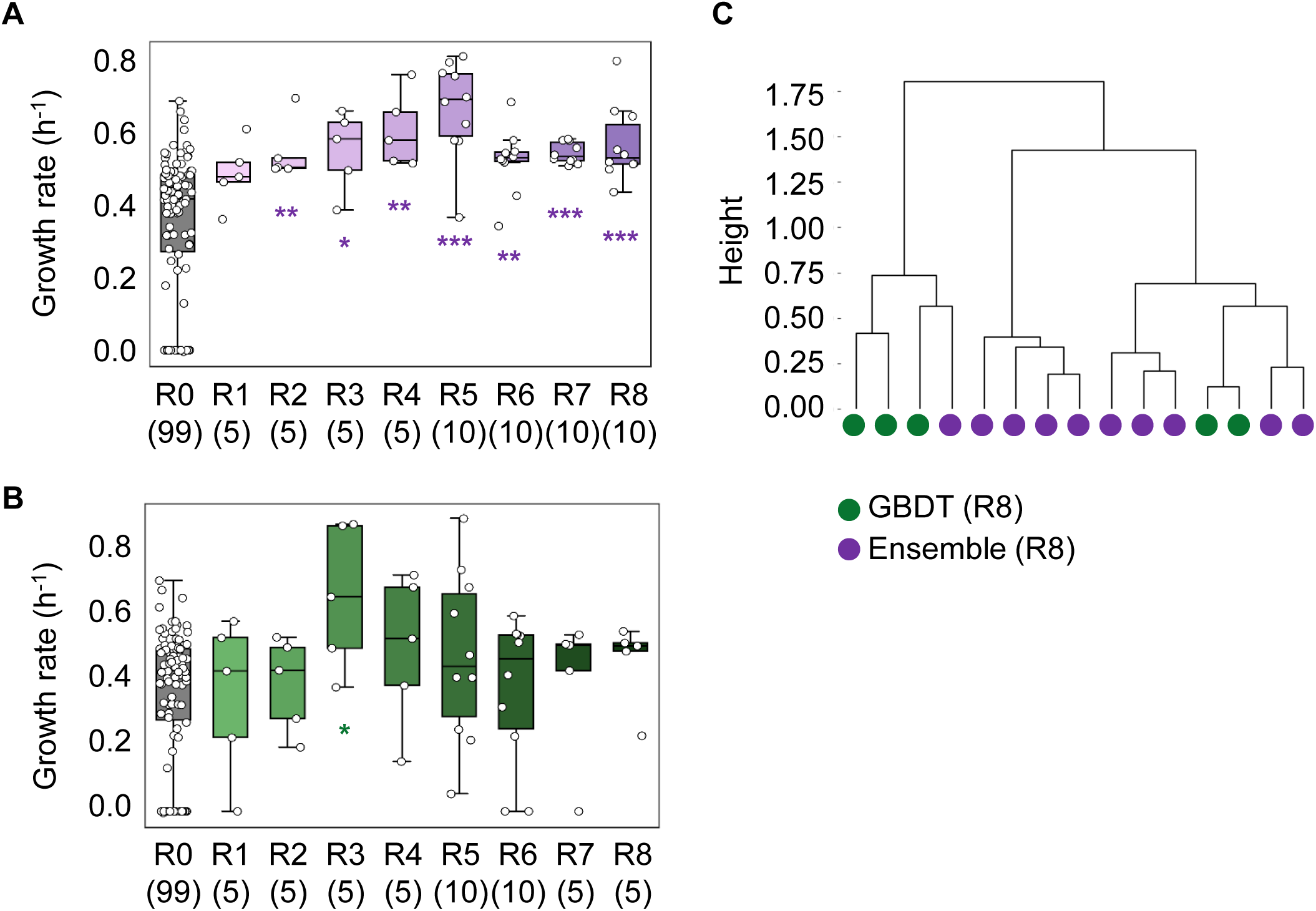
ML-guided medium fine-tuning. **A.** Experimentally validated growth rates obtained during active learning with the Ensemble model. **B.** Experimentally validated growth rates obtained during active learning with the GBDT model. Open circles represent the mean growth rates of biological replication. Asterisks indicate the statistical significance as follows: *, p<0.05; **, p<0.01; ***, p<0.001. **C.** Hierarchical clustering of the medium compositions predicted in R8. The ward method was used. Green and purple indicate the media predicted by the GBDT and Ensemble models, respectively.

### ML model-dependent manner of habitat searching for fitness increase

The patterns of fine-tuned medium compositions were differentiated between the two ML models. The active learning stopped at R8 due to saturated growth rates. Clustering of the medium compositions predicted at the final round (R8) revealed that GBDT and Ensemble guided the differentiated patterns of fitted media for Anc (Fig. 2C), independent of the clustering methods (Fig. S4). Such differentiated patterns of medium fine-tuning were likely initiated in the earlier rounds, e.g., R5 (Fig. S5). The distinct directional fine-tuning of medium compositions was further verified by analyzing all 110 media that were both predicted and tested in active learning. The results revealed the diverse directions of the two ML methods in guiding medium fine-tuning (Fig. 3). Significant positive and negative correlations were identified between two main principal components (PC1 and PC2) of the medium compositions predicted by the Ensemble and GBDT models, respectively (Fig. 3A). However, no significant correlation was observed between PC1 and PC2 when PCA was conducted to the medium compositions predicted by the two ML methods separately (Fig. S6). These findings indicated the directions (e.g., rules, mechanisms) of medium fine-tuning were distinct through GBDT and Ensemble, although both ML models fine-tuned the medium compositions randomly.

**Figure 3.**
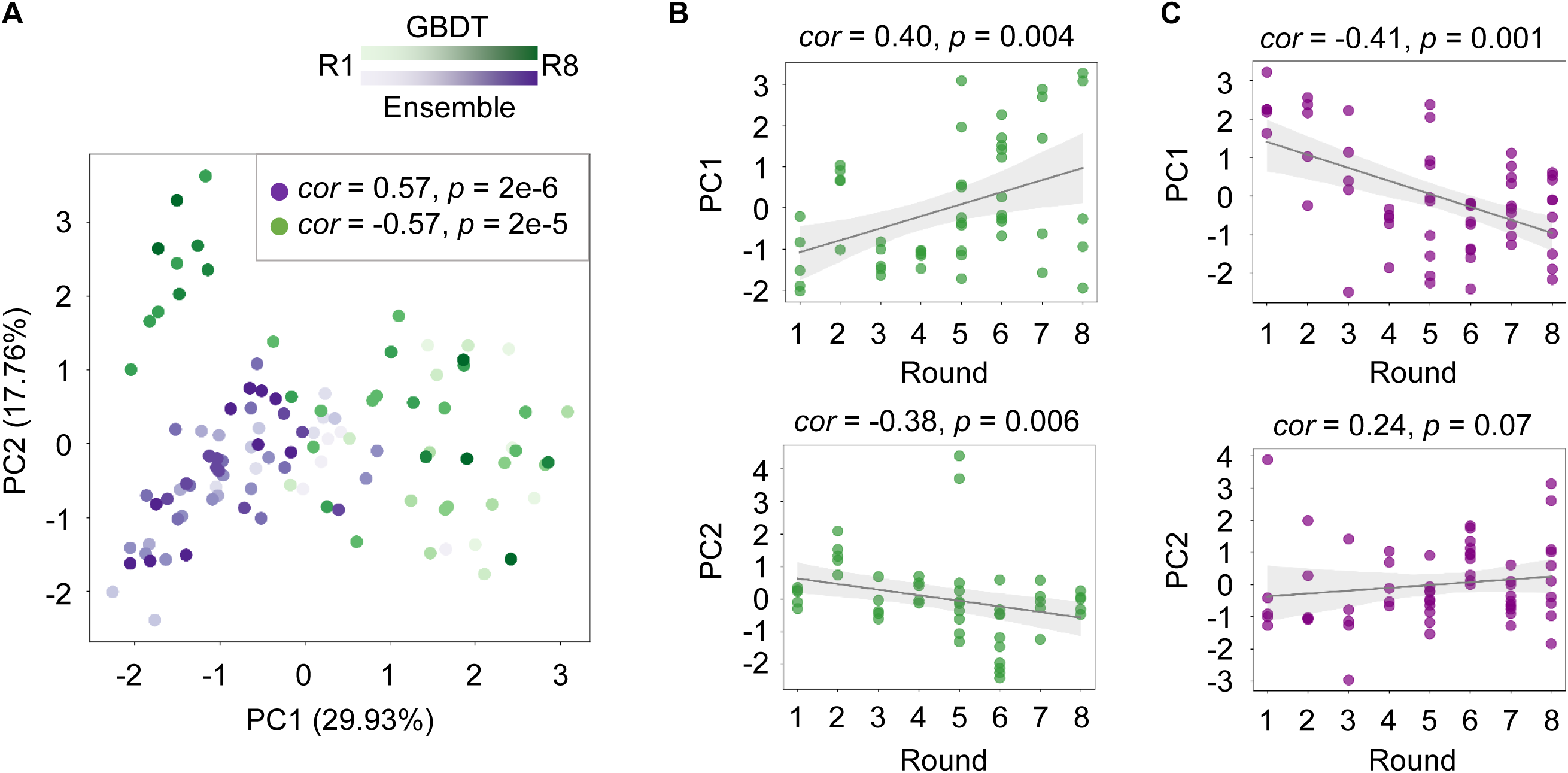
PCA of medium compositions. **A.** PCA of 110 tested media from R1 to R8. Green and purple indicate the media predicted by the GBDT and Ensemble models, respectively. Color gradation represents the varied rounds (R1∼R8) of active learning. Spearman’s correlation coefficients and *p*-values are indicated. **B.** Relation between PCs and active learning rounds in the GBDT model. **C.** Relation between PCs and active learning rounds in the Ensemble model. Upper and bottom panels show PC1 and PC2, respectively. Five to ten tested media are indicated as dots. Spearman’s correlation coefficients, p-values, and 95% confidence intervals are indicated.

The active learning round proceeding was associated with directional fine-tuning patterns. The positive and negative correlations of the eight GBDT rounds were significant to PC1 and PC2, respectively (Fig. 3B). A negative correlation was detected between PC1 and the eight Ensemble rounds (Fig. 3C), which suggested a gradual directional search for the better-fitted habitats guided by ML. It agreed with the finding that the PC1-PC2 correlations are significant from R1 to R8 (Fig. 3A) but are insignificant from R1 to R5 (Fig. S7). The ML models guided directional medium fine-tuning (habitat searching) to approach increased fitness. Although it’s unknown whether such directional habitat searching occurs in nature, it demonstrated the successful application of ML in experimental ecology to identify growth environments that are better suited for the target bacterium across numerous possibilities.

### Comparing fitted habitats to fitted genomes of equivalent fitness increases

Fitted genomes obtained by experimental evolution were selected from our previous study ^31^. Six fine-tuned media (OPTs) that showed a significant increase in growth fitness (Fig. 4A, blue) and five evolved lineages (EVOs) that exhibited increased growth rates (Fig. 4A, magenta) were subjected to transcriptome analysis. Both OPTs and EVOs exhibited equivalent growth rates and growth change in comparison to Ori (Fig. 4B), suggesting the changes in genomes and in habitat led to a comparable impact on growth fitness. The genome mutations fixed in the five EVOs varied from two to 13, compared to Anc (Fig. 4C), reflecting the genetic diversity of fitted genomes generated by the evolutionary strategy. The compositions of the six OPTs varied in two to all eight components, compared to each other and M63 (Fig. 4D), indicating the diversity of fitted habitats resulting from the ecological strategy.

**Figure 4.**
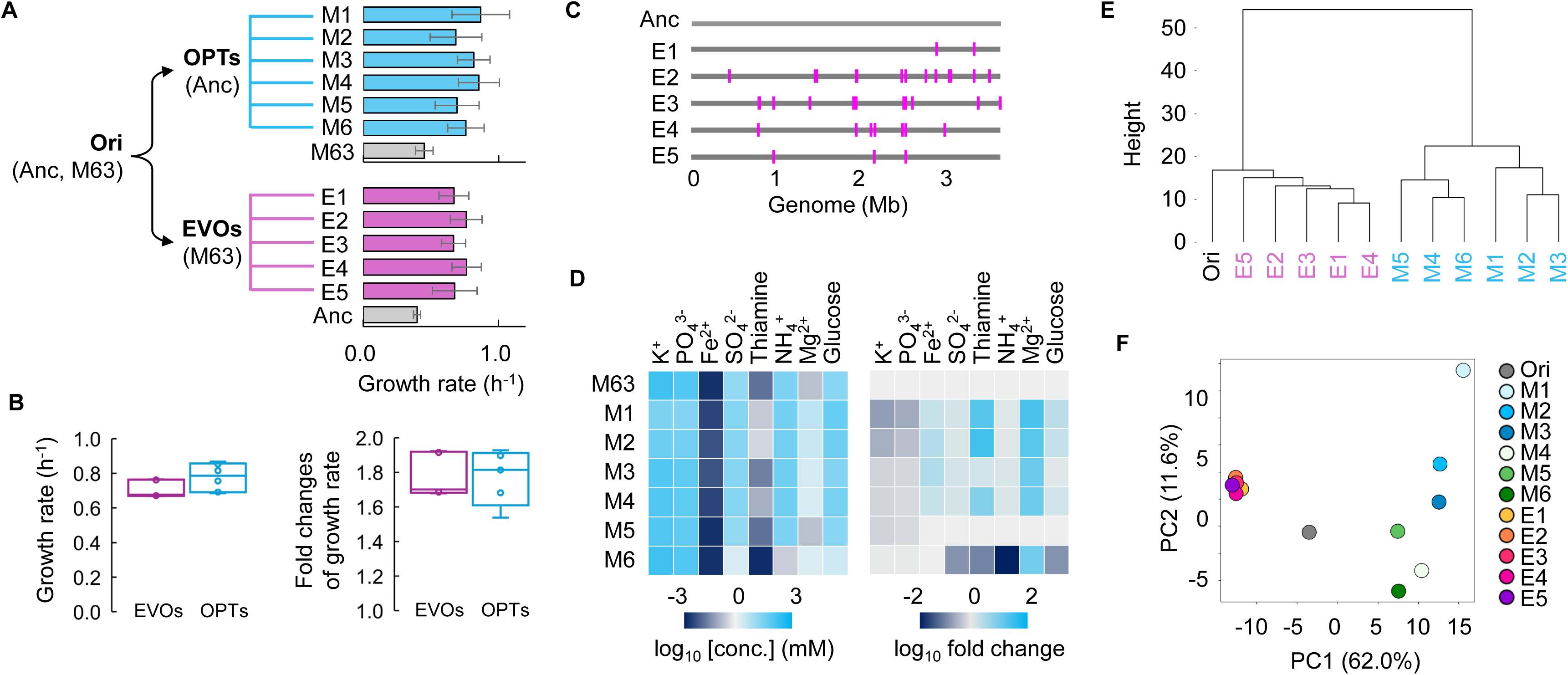
Comparison of fitted genomes and fitted habitats. **A.** Selected OPTs and EVOs. Six OPTs (M1∼M6) and five EVOs (E1∼E5) are used for comparison. The growth rates of Anc grown in OPTs and EVOs in M63 are shown, and the standard errors of biological replicates (n = 6) are indicated. **B.** Mean growth fitness of OPTs and EVOs. Left and right panels indicate the mean fitness and mean fitness changes, respectively. Megenta and blue represent the means of six OPTs and five EVOs, respectively. **C.** Genetic mutations accumulated in EVOs. Genomic positions of mutations fixed during experimental evolution are highlighted in magenta on the linear genomes. **D.** Medium compositions of OPTs. Eight chemical components that are fine-tuned for habitat reconstruction are indicated. Color gradations represent the gradients of chemical concentrations and fold changes on a logarithmic scale. **E.** Hierarchical clustering of transcriptomes. The Ward method is used. Individual OPTs and EVOs are indicated. **F.** PCA of transcriptomes. Color variation indicates individual OPTs and EVOs.

Transcriptome analysis of EVOs and OPTs was conducted to investigate how evolutionary and ecological strategies reorganize the gene expression for fitness increase. Clustering analysis revealed that the genome-wide expression patterns were divergent between EVOs and OPTs (Fig. 4E, Fig. S8), indicating that evolutionary and ecological strategies led to the differentiation of transcriptome reorganization. PCA further observed that the transcriptome reorganization of EVOs directed to a highly similar expression pattern, whereas that of OPTs offered highly varied expression patterns (Fig. 4F). Both evolutionary and ecological strategies succeeded in finding multiple solutions for fitness increase by achieving the fitted genomes and media, respectively, however, they reorganized the transcriptomes apart from that of Ori in a distinct manner. EVOs tended to a similar directional transcriptome reorganization, while OPTs provided a variable degree of transcriptome reorganization, despite EVOs and OPTs causing equivalent fitness improvements.

### Different reorganization manners and common metabolisms in transcriptomes between fitted habitats and fitted genomes

The transcriptome changes caused by medium fine-tuning presented different patterns from the viewpoints of individual genes. The abundance of differentially expressed genes (DEGs) varied significantly among the six OPTs, ranging from 315 to 1,018 genes, which enriched one to ten gene orthologs (GOs) (Fig. 5A). Note that no mutated genes were identified in the DEGs. Only 70 DEGs overlapped in common, and none overlapped in enriched GOs. Contrastingly, the DEGs identified in EVOs were highly abundant, ranging from 1,215 to 1,598 genes, which shared 718 universal DEGs (Fig. 5B). Every 12 ∼ 19 GOs were enriched in the five EVOs (Fig. 5B), of which 11 overlapped (Fig. S9). The numbers of both DEGs and the enriched GOs varied in OPTs but were similar in EVOs. 22 GOs were enriched in the 718 common DEGs of EVOs, and no GO was significantly enriched in the 70 common DEGs of OPTs. Twenty-six genes overlapped between 70 OPTs- and 718 EVOs-derived DEGs (Fig. 5C), in which three GOs relating to arginine and glutamine processes were significantly enriched (Fig. 5D). These differences between OPTs and EVOs were consistent with their divergent gene expression patterns (Fig. 4E-F). The equivalent fitness increase achieved by OPTs and EVOs was commonly attributed to several genes participating in arginine and glutamine metabolism, which was unreasonable, although it’s a known pH-buffering and nitrogen storage pathway ^40,41^.

**Figure 5.**
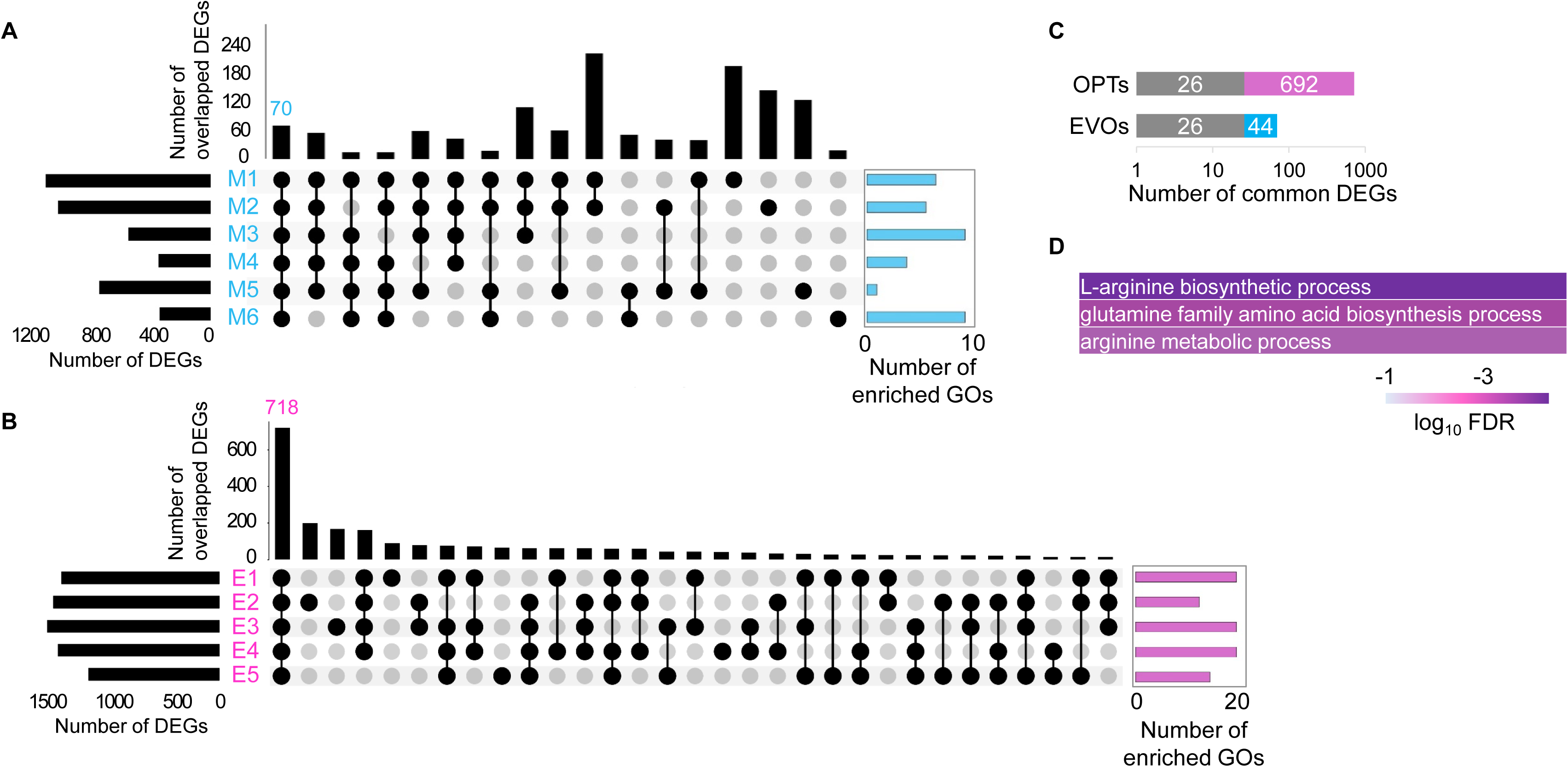
DEGs determination and GO enrichment. **A.** DEGs identified in OPTs. Black and blue bars represent identified DEGs and enriched GOs, respectively. Filled circles connected with black lines indicate the overlapped DEGs among OPTs. The number of common DEGs across all OPTs is indicated. **B.** DEGs identified in EVOs. Black and magenta bars represent identified DEGs and enriched GOs, respectively. Filled circles connected with black lines indicate the overlapped DEGs among EVOs. The number of common DEGs across all EVOs is indicated. **C.** Number of the overlapped and distinct DEGs between OPTs and EVOs. **D.** Enriched GOs. Color gradation represents FDR on a logarithmic scale.

Alternatively, analyzing transcriptome changes from the viewpoints of functional gene networks observed universal changes caused by EVOs and OPTs. Gene set enrichment analysis (GSEA) was performed to evaluate the significance of the changes in the entire gene sets (all genes that participated in the biological process). Five out of 15 gene sets were commonly identified in OPTs (Fig. 6A), compared to ten out of 12 enriched gene sets in common in EVOs (Fig. 6B). These gene sets were different from those enriched in the 26 DEGs, indicating the functional reorganization for fitness increase at the level of gene network (pathway, process, etc.) but not specific genetic function. Intriguingly, the five common gene sets in OPTs were all included in the ten common gene sets in EVOs, suggesting they were the common biological processes responsible for fitness increase, regardless of the genetic or environmental causes. These metabolisms might play a universal role in fitness increase caused by either genetic or environmental adaptation. Such a profound commonality in evolutionary and ecological strategies for fitness increase indicated a general eco-evolutionary mechanism underlying the transcriptome reorganization for fitness increase.

**Figure 6.**
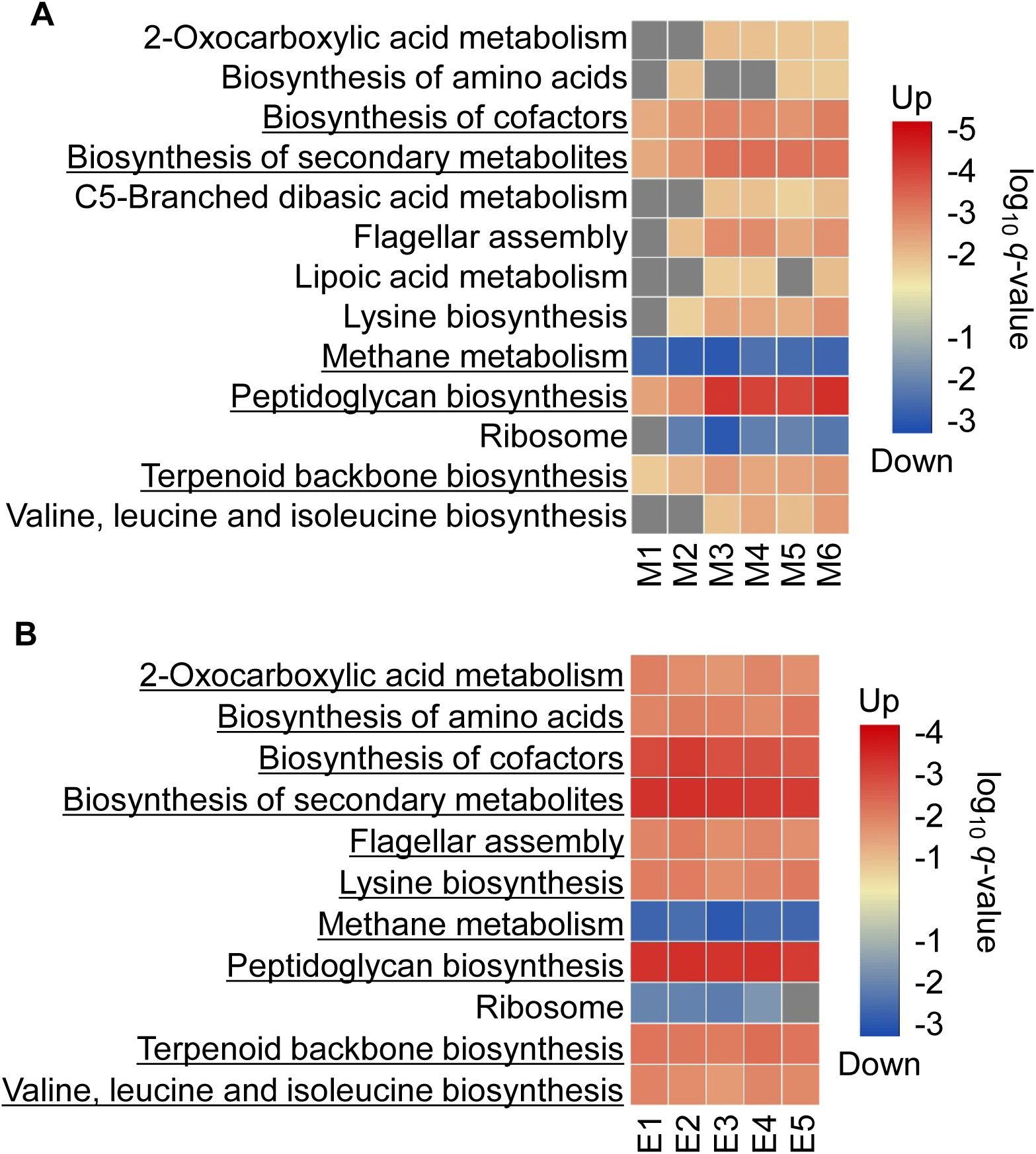
GSEA of EVOs and OPTs. **A.** GSEA of OPTs. **B.** GSEA of EVOs. Color gradation indicates the statistical significance, *q*-value, on a logarithmic scale. Red and blue color tones reveal transcriptional activation and inhibition, respectively.

### ML-navigated habitat reconstruction might avert genetic restriction cleared by evolution

Whether any compensation or replacement in the biological process could occur between fitted genomes and habitats was further investigated. ML model-dependent contributions of medium components to fitness increase were evaluated. Converting the medium compositions to their ionic formats resulted in eight chemical components, of which the impacts (i.e., feature importance) on growth rates were predicted (Fig. 7A). A single component, i.e., K^+^, presented the highest feature importance in the GBDT-navigated media, indicating its high priority in deciding the growth rate. Alternatively, multiple components, e.g., SO_4_^2-^, Fe^2+^, and thiamine, shared relatively high feature importance in Ensemble-navigated media (Fig. 7A). This suggested that the GBDT and Ensemble models guided the determinative and cooperative working principles of medium components to increase the growth fitness, respectively. The patterns of the feature importance were significantly divergent between the two ML models, as well as when compared with the initial pattern (Fig. 7B). This indicated that the ML-guided medium fine-tuning targeted varied chemical properties, nutritional functions, and biological pathways. Such differentiation in working principles was evident throughout the entire active learning process. The determinative and cooperative working manners of medium components impacted growth were observed at earlier rounds, e.g., R5 (Fig. S10), which agreed with the clustering of medium compositions (Fig. S5).

**Figure 7.**
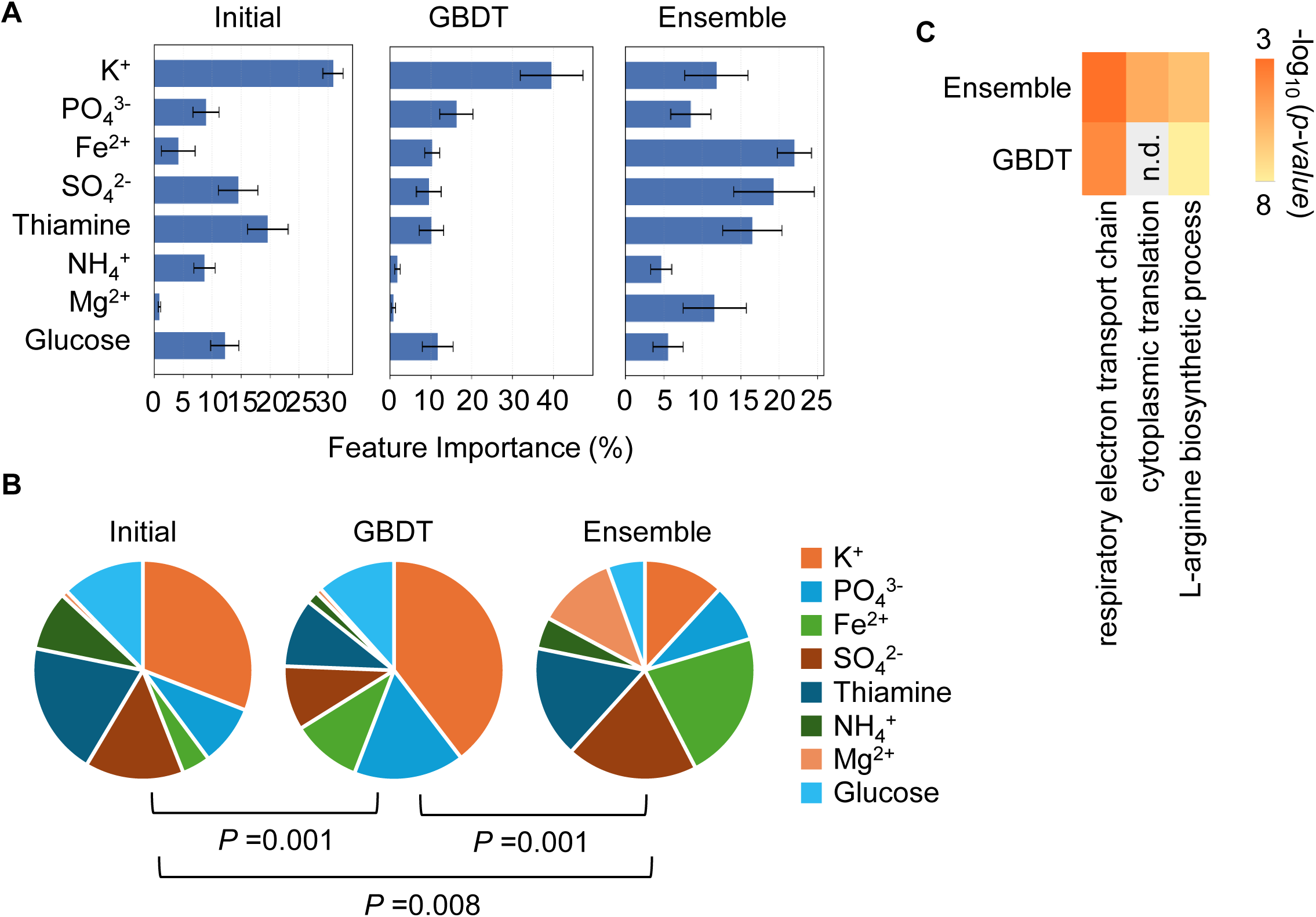
Contributions of medium components to fitness increase. **A.** Bar graphs of feature importance. Standard Errors indicate five repeated predictions. **B.** Pie graphs of feature importance. Initial, GBDT, and Ensemble indicate the predictions using the initial 99 media, the 50 GBDT-navigated media, and the 60 Ensemble-navigated media, respectively. Statistical significance of the pattern comparison is indicated (PERMANOVA). **C.** Enriched GOs in the DEGs identified in the GBDT- and Ensemble-predicted OPTs. Color gradation represents the statistical significance of enrichment, *p*-value, on a logarithmic scale.

Such an ML-guided habitat reconstruction might fine-tune metabolism or regulation, bypassing genetic bottlenecks for fitness increase. Representatively, most EVOs accumulated mutations in the genes of *cysP*, *cysU*, and *cysA*, of which the products are the sulfate/thiosulfate transporter subunits, agreeing with the functional enrichment of DEGs enriched in sulfur compound biosynthetic and sulfur compound metabolic processes. This evolutionary strategy was somehow compensated by the ecological strategy targeting the sulfate ions, significantly in the Ensemble-guided medium fine-tuning (Fig. 7). The functional enrichment of DEGs in OPTs failed to detect any sulfur-related processes, suggesting that the fitted habitats (media) avoided disturbing the biological process essential for fitness increase. Besides, gene mutations in other transporters were also found in EVOs, which might be compensated by the OPTs targeting potassium ions, particularly in GBDT (Fig. 7A). Consistently, the respiratory electron transport chain, a process participated by potassium ions ^42^, was significantly enriched in the DEGs of OPTs (Fig. 7C). These findings suggested that ML-navigated ecological strategy successfully conquered the genetic bottlenecks required for mutations in sulfur and other transport.

## Discussion

Both experimental evolution and experimental ecology achieved fitness increases by finding a variety of mutated genomes and fine-tuned media, respectively. Although both strategies led to equivalent improvements in growth fitness, the changes that occurred among the multiple solutions were varied (Fig. 8A). EVOs exhibited more DEGs and enriched functions of high overlaps across the five fitted genomes, whereas OPTs showed fewer DEGs and functions of few overlaps. It indicated that the evolutionary and ecological strategies adopted different working manners to increase growth fitness, i.e., genetic changes for generality and habitat changes for diversity. Nevertheless, a few surrounding biosynthetic processes, e.g., cofactors, secondary metabolites, peptidoglycan, etc., were commonly reorganized (Fig. 6), indicating a general and obligatory mechanism for fitness increase.

**Figure 8.**
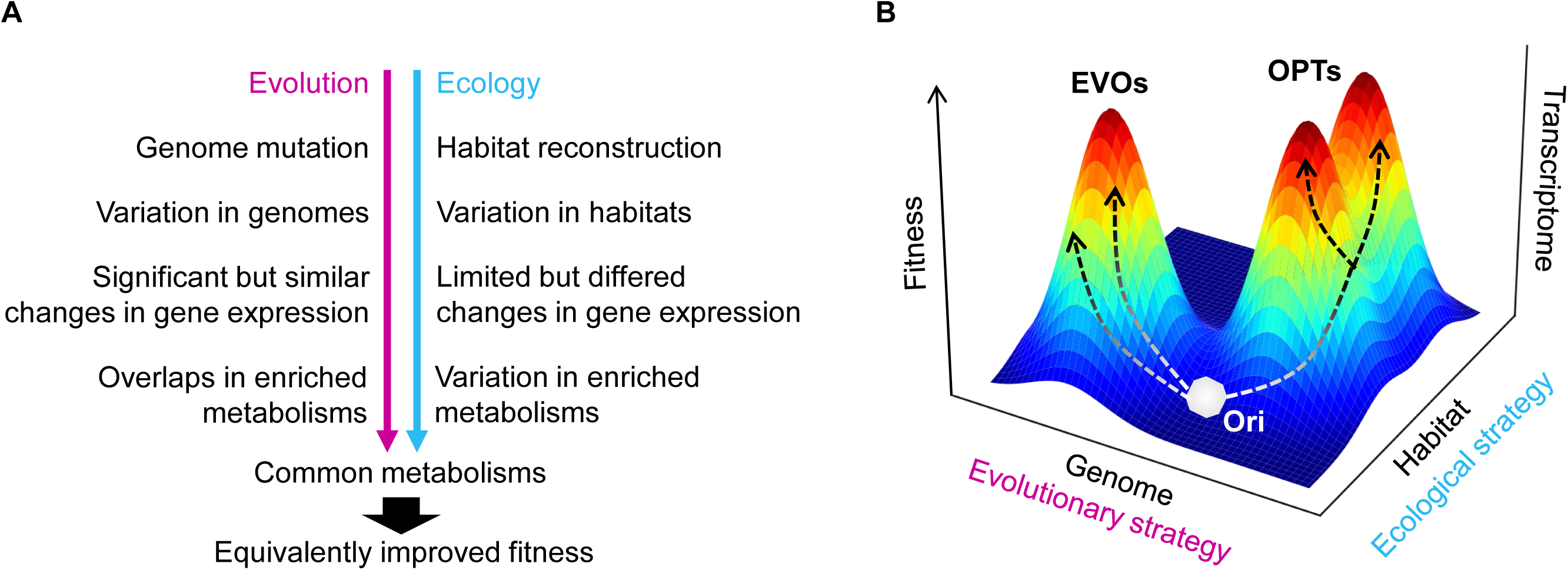
Overview of evolutionary and ecological strategies for fitness increase. **A.** Summary of the findings in the present study. **B.** Fitness landscape on an evolutionary and ecological space. The evolutionary and ecological paths for Ori to approach higher fitness are shown in arrowed broken lines.

These experimental demonstrations revealed that habitat reconstruction was not simply a replacement for genetic improvement but also a computationally guided complementary path to discover fitness landscapes (Fig. 8B). Habitat reconstruction adjusted substrate availability (i.e., concentrations of medium components) advantageous for refined metabolic pathways; comparably, genome evolution adjusted gene function (i.e., mutations) to benefit the improved metabolism. ML-guided fine-tuning did not change the chemical components but their balance in the habitat, which differed from what was generally studied by adding, removing, or replacing the substrates in the growth media ^43,44^. Diverse ML algorithms drove distinguished paths to climb the fitness mountains, compared to the serial transfer leading to an identical path for fitness increase. It’s unclear whether the variation in the methodology of serial transfer could cause multiple paths, as ML provided. Although the evolutionary and ecological strategies approached distinct paths, they ultimately acted on transcriptomes and achieved comparable fitness benefits (Fig. 8B). This finding strongly suggested there were multiple routes of ecological niche construction to escape evolutionary bottlenecks for fitness increase, supporting the bypass mechanism for resistance ^45^.

The present study allowed us to have some intriguing insights. Among the six OPTs and five EVOs, the central machinery, the ribosome, was not commonly enriched (Fig. 6), indicating that fine-tuning the translation was not secure for fitness increase, regardless of evolutionary or ecological strategies. It was out of expectation, as extensive studies reported the contribution of ribosomes to bacterial growth in a quantitative correlated manner ^46–48^. Instead, five metabolic processes universally enriched in both EVOs and OPTs were somehow reasonable but remained largely mysterious. For instance, the cofactor biosynthesis might relate to the contribution of Fe^2+^ in OPTs and the mutated *nadR* and *acpP* in EVOs, according to their molecular mechanisms ^49,50^. The methane metabolism, commonly down-regulated in OPTs and EVOs, failed to be explained, indicating unknown mechanisms that require addressing. In addition, the six OPTs generated through active learning not only increased the growth rates by ∼30% but also highlighted ML model-specific ecological strategies. For instance, the Ensemble model prioritized sulfate (SO₄²⁻) and iron (Fe²⁺) ions, possibly due to their participation in organic carbon utilization, fermentation, and respiration ^51^, affecting bacterial growth.

As a challenge of experimental ecology by active learning, two different ML models successfully identified the fitted media, and the differences could be discerned. Out of the six OPTs, M1-M4 and M5-M6 were derived from the Ensemble and GBDT models, respectively. Due to the ML-oriented medium search, distinct mechanisms for adapted habitats were identified. GBDT drove a highly correlated change in medium components (Fig. 3B) upon the initial habitats (pool of media), which maintained the working principle of a single key component determining the growth fitness (Fig. 7A-B). The Ensemble method revealed partially correlated changes (Fig. 3C), which triggered a distinct working principle of the corporation of multiple components in growth decision-making (Fig. 7A-B). Although both working principles led to an equal increase in fitness, the transcriptome changes presented different patterns. For instance, the enriched gene sets showed greater dissimilarity among the OPTs derived from Ensemble (Fig. 6A). Additional functional enrichment analyses, e.g., gene category, regulon, etc., revealed diverse trends between the OPTs derived from GBDT and those derived from Ensemble (Fig. S11). The results strongly suggested that employing varied ML models could allow us to discover various working principles of fitted habitats.

The limitations of using ML in this study were both the ML models and the datasets. Whether any other ML models would result in alternative findings remained unclear. The algorithms from a computational or statistical viewpoint could cause distinct outcomes derived from different ML models ^52–54^. The Ensemble model was somewhat better than GBDT in prediction (Fig. 2, Fig. S2, Fig. S3). Aggregating multiple algorithms in the Ensemble model might reduce the risk of overfitting ^52,53^, leading to a highly accurate and robust prediction, particularly for biological datasets ^53^. In addition, the size of the initial training data used here was 99 medium variations of the eight elements. Whether the 99 variation was reasonable was unknown. A considerable variation of medium compositions was initially tested to mimic a wide range of ecological habitats. Achieving the medium compositions that showed improved growth fitness by this approach was a chance occurrence. In summary, employing ML to search for the fitted habitats provided a successful demonstration of ecological searching in the laboratory and decoupled fitness increases from genetic change, allowing us to investigate the relative contributions of ecological vs. evolutionary mechanisms. It revealed how environmental change could precede fitness increase and offered a methodological platform for non-genetic adaptation in biotechnology and microbiome engineering.

## Materials and Methods

### Bacterial strain and stocks

The *E. coli* strain was from the National BioResource Project, National Institute of Genetics, Japan. It was derived from the wild-type strain W3110, described previously ^37,55^. The bacterial stocks were prepared for the growth assay to reduce the experimental errors caused by the initial bacterial inoculation, as described previously ^16,56^. The bacterial strain was cultured in a test tube containing 5 mL of M63 basal medium and incubated in a biological shaker (TAITEC) at 200 rpm and 37 °C. The cell culture was terminated at the end of the exponential phase, aliquoted into 750 μL portions in 1.5 mL microtubes (Watson), and mixed with 250 μL of 60% glycerol solution. Subsequently, 50 μL of the mixed solution was added to each new 1.5 mL microtube and stored at −80℃. Multiple aliquots (original culture solution) were prepared in one batch and used all at once to avoid deviations caused by repeated freeze-thaw cycles or pre-cultivation. The aliquots were used only once, and any remaining culture solution was discarded.

### Medium preparation

All media were prepared using solution stocks of pure chemical compounds, including KH_2_PO_4_, K_2_HPO_4_, FeSO_4_, Thiamine, (NH_4_)_2_SO_4_, MgSO_4_, and glucose, which were purchased from Sigma or Wako. These compounds were dissolved in pure water (MilliQ), yielding corresponding concentrations of 0.385, 0.615, 0.0018, 0.015, 3, 0.2, and 1 mol/L, and the stock solutions were prepared as described previously ^16,30^. These solutions were then sterilized by filtration using a syringe with a 0.22 μm pore size (hydrophilic PVDF membrane, Merck) or by autoclaving at 121 °C for 20 minutes (TOMY). The sterilized chemical solutions were divided into 1000 μL portions and stored in 1.5 mL microtubes (Watson). The solution stocks were stored at −30°C for future use. The details of the solution stocks are summarized in Table S1. The media used for the growth assay were prepared by mixing these solution stocks just before bacterial culture. The solution stocks were single-use, and the remaining ones were discarded to avoid repeated freezing and dissolution.

### Growth assay and calculation

The bacterial stocks were used for the high-throughput growth assay in a 200 μL culture, as performed in our previous studies ^16,30^. In brief, 96-well microplates (Coaster) were used for bacterial culture and incubated at 37°C in a plate reader (EPOCH2, BioTek), with shaking at 567 cycles per minute (cpm) for 48 hours. The temporal changes in OD600 were measured at 30-minute intervals. 6 to 42 biological replicates were conducted for each medium. A total of 690 assays were performed. The OD600 records were exported from the plate reader and processed using Python, as previously described ^16,35^. The growth rates were calculated accordingly.

### Machine learning

Machine learning was performed using Python 3.11.7 programming environment ^57^. The GBDT model utilized the gradient boosting decision tree (GBDT) for both model construction and prediction due to its interpretability, as demonstrated in our previous study ^39,56^. The Ensemble model constructed multiple basic regressors, including Support Vector Regression (SVR), Gradient Boosting Decision Trees (GBDT), Multi-Layer Perceptrons (MLP), and K-Nearest Neighbor Regression (KNN). Grid search (GridSearchCV) was used to tune the hyperparameters of each submodel. These submodels were then integrated into a StackingRegressor, with a linear regressor as the final meta-learner. A multi-layer feedforward neural network was constructed using Keras as part of the ensemble model. The network consists of multiple fully connected layers with ReLU activations, using the Adam optimizer and mean squared error as the loss function. To infer the optimal input variable combinations from the model, a genetic algorithm implemented using the DEAP library was employed. An adaptive fitness function is defined, which predicts the phenotypic performance of potential input combinations using the trained stacked regressor model. Optimization search employs strategies such as roulette wheel selection, single-point crossover, and floating-point mutation. The optimization objective is to maximize the predicted target variable (e.g., maximum specific growth rate). The optimal individuals were selected from each generation of optimization, decoded back to the real space, and their phenotypic performance was predicted. These are then incorporated into the final dataset for subsequent analysis or model iteration. The code used for the two ML models is available on G*itHub* (see Code Availability).

### Active learning

As shown previously, the GBDT and Ensemble algorithms were utilized for active learning, as performed previously ^34,36^. Model construction and prediction were performed using Python. The explanatory and objective variables were medium composition and MaxRate, respectively. They were evaluated in data processing using MaxRate values from cultures grown in the M63 medium. Initially, 690 growth curves were used as training data for model construction and data processing. Every five to ten medium compositions predicted to exhibit the highest growth rates were subjected to experimental validation in each round. The experimental results were incorporated into the training data for the subsequent round of learning, model construction, prediction, and validation. The rounds were repeated eight times (R1-R8), and the media combinations that substantially increased the MaxRate values were selected from them.

### Evaluation of ML models

To evaluate the predictive performance of the regression model at the sample level, the mean squared error (MSE) was used as the evaluation metric. We used the mean_squared_error function from the scikit-learn library to calculate the squared difference between the predicted values and the actual values. This function provides a quantitative measure of prediction bias, i.e., the lower the MSE value, the higher the prediction accuracy. By calculating the MSE for all samples, this method helps identify cases with poor prediction performance and supports subsequent model optimization.

### Feature importance

The log-transformed concentrations of the components were used for GBDT prediction. The GBDT algorithm was used to perform a variable importance ranking analysis, as conducted in our previous studies ^16,56^. The train_test_split function was used to randomly split the dataset into a training set and a test set (in an 8:2 ratio). To improve model stability and reduce bias caused by random splitting, the model was trained five times, and feature importance scores were extracted after each run and standardized to percentage form. Subsequently, the average of the feature importance scores obtained from the five model training runs was taken as the final importance assessment for each feature.

### RNA sequencing

RNA sequencing was performed as described previously ^15,31^. Bacterial culture was performed in a test tube at a 5 mL volume and was incubated in a bioshaker (BR-23FP, Taitec) at 200 rpm, 37°C. A precision particle size analyzer (Multisizer 4, Beckman Coulter) with an aperture of a pore size of 20 μL was used to evaluate the cell concentration. 20 μL of culture was suspended in a dedicated 25-mL cuvette (Beckman Coulter) containing 10 mL of diluent (Isoton II, Beckman Coulter). The *E. coli* cells were collected during the exponential growth phase (i.e., 5 × 10^7^ ∼ 2 × 10^8^ cells/mL). The test tubes were quickly transferred to ice and mixed with Falcon containing 5 mL of a 10% phenol-ethanol solution. The Falcon was subsequently centrifuged at 7,000 rpm, 4°C, for 3 min, the supernatant was removed, and the pellet was frozen at −80°C for future use. The pellets were thawed for at least 10 minutes, and the total RNA was purified using the RNeasy Mini Kit (QIAGEN) and the RNase-Free DNase Set (QIAGEN) according to the manufacturer’s instructions. Purified total RNA was dissolved in RNase-free water and frozen at −80°C. Whole genome sequencing was performed commercially by Azenta Life Sciences Japan, headquartered in Burlington, USA. Biological replications were performed for all conditions (N = 2∼5).

### RNAseq data processing and normalization

The reference genome W3110, obtained from GenBank under accession number AP009048.1, was mapped to the paired-end FASTQ obtained by RNA-seq, as previously described ^15,31^. The RNAseq datasets of OPTs were deposited in the DDBJ Sequence Read Archive under the accession numbers DRA017794 and DRA019044. The raw datasets of EVOs were acquired from the DDBJ Sequence Read Archive under the accession number DRA013662. A total of 3,290 genes were used for the transcriptome analyses. The obtained read counts were converted to FPKM values according to the gene length and total read count values. Global normalization of the FPKM values was performed to achieve a uniform mean value on the logarithmic scale across all datasets. The gene expression level was determined as the logarithm of the FPKM value. The dataset was used for the transcriptome analyses.

### Transcriptome analyses

Transcriptome analyses were performed using R (version 4.3.0)^58^, as described in our previous studies ^15,31^. Hierarchical clustering and principal component analysis (PCA) were performed using “clusterSample” and “prcomp”, respectively. In clustering analysis, the “dist.method” was set as “spearman” and the “hclust.method” was set as “ward.D2”, “complete”, and “average” for multiple evaluations. In PCA, the “scale” was set as “F.” Differentially expressed genes (DEGs) were identified using the R package DESeq2 ^59^. The read counts and logarithmic FPKM values were used as the input data for DESeq2. The differentially expressed genes (DEGs) were identified based on the false discovery rate (FDR). Functional enrichments of DEGs were performed based on transcriptional regulation (Regulon) ^60^, gene categories (GCs) ^61^, gene ontologies (GOs) ^62,63^, and metabolic pathways ^50^. A total of 53 transcriptional factors (TFs) comprising more than 10 regulatees and 19 GCs of more than 30 genes were subjected to enrichment analysis. A total of 20 GCs were used in the analysis. The statistical significance of the DEGs enriched in TFs and GCs was evaluated using the binomial test with Bonferroni correction. The functional enrichment of GOs was performed using the Database for Annotation Visualization and Integrated Discovery (DAVID) version 2021. The statistical significance was determined according to FDR.

### Gene set enrichment analysis

Gene set enrichment analysis (GSEA) ^64,65^, identifying the significant enrichment of predefined gene sets within the ranked list of DEGs, was performed as previously described ^66^. All analyses were performed in the R environment (version 4.3.0), primarily using R packages such as org.EcK12.eg.db, clusterProfiler, pathview, and enrichplot. First, the differentially expressed gene symbols and their log2FC values obtained from preprocessing were converted to NCBI ENTREZ IDs and then sorted in descending order of log2FC values to construct an ordered gene list. Subsequently, they were further converted to KEGG IDs, and a named vector with log2 fold change (log2FC) values was constructed. Pathway enrichment analysis was performed using the gseKEGG function (significance threshold *p* < 0.05).

## Data availability

The RNAseq data obtained in the present study are available at https://ddbj.nig.ac.jp/public/ddbj_database/dra/fastq/DRA017/DRA017794/ and https://ddbj.nig.ac.jp/public/ddbj_database/dra/fastq/DRA019/DRA019044/

## Code availability

The ML code used in the present study is available at https://github.com/LuZipeng/ML.

## Author contributions

ZL conducted the experiments, data analyses, and drafted the manuscript. BWY conceived the research, validated the analyses, and rewrote the manuscript. All authors approved the final manuscript.

## Acknowledgments

We thank NBRP (Japan) for providing the *E. coli* strain. This work was supported by the JSPS KAKENHI grant number 25K02259 (to BWY).

