## Supplemental figures and table for "Equivalent fitness increase achieved by active learning-navigated habitat reconstruction and evolution-induced genome mutation"

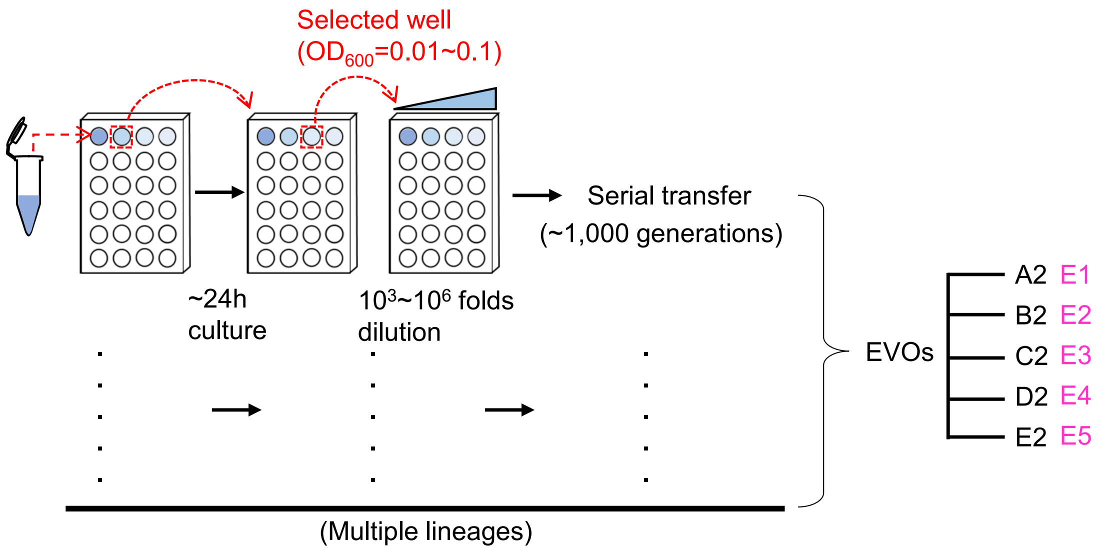


**Figure S1 Experimental evolution.** Serial transfer was performed in the previous study (see Materials and Methods) to obtain the evolved populations of increased fitness. Five evolved lineages of the most significant growth increase are indicated. The labels used in the previous and present studies are indicated in black and magenta, respectively.

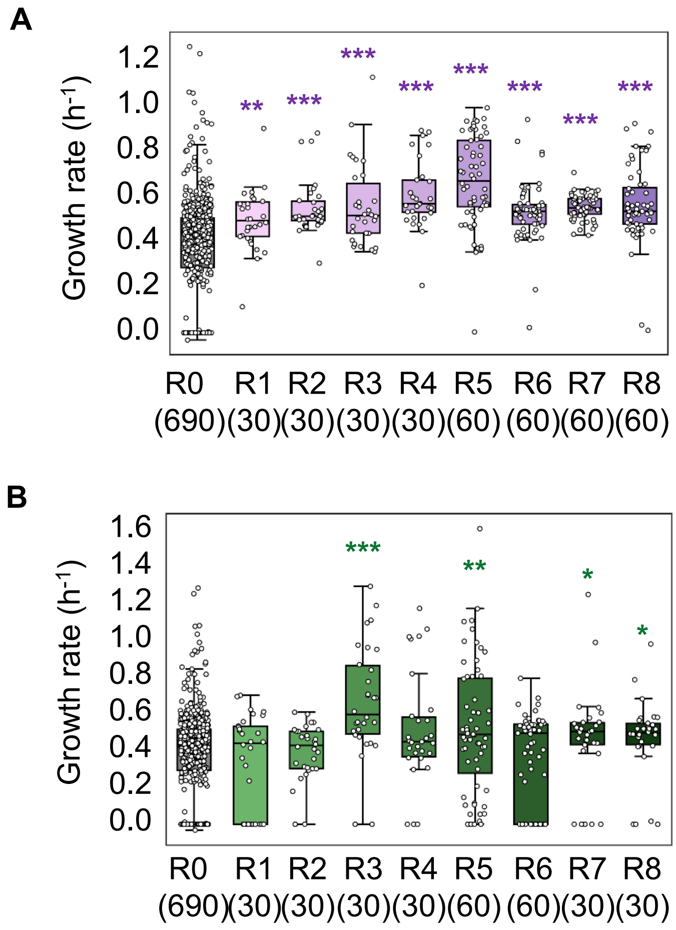


**Figure S2 ML-guided medium fine-tuning. A.** Experimentally validated growth rates obtained during active learning with the Ensemble model. **B.** Experimentally validated growth rates obtained during active learning with the GBDT model. Open circles represent individual growth rates acquired per well (assay). Asterisks indicate the statistical significance as follows: *, p<0.05; **, p<0.01; ***, p<0.001.

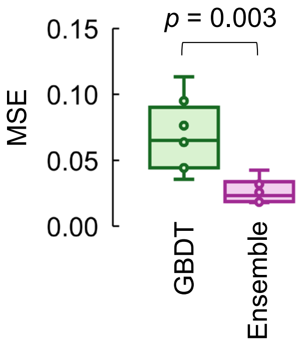


**Figure S3 Prediction accuracy of ML models.** The prediction accuracy across all eight rounds of active learning is evaluated by the Mean Squared Error between the predicted growth rate and the experimentally validated growth rate. Statistical significance of the t-test is indicated.

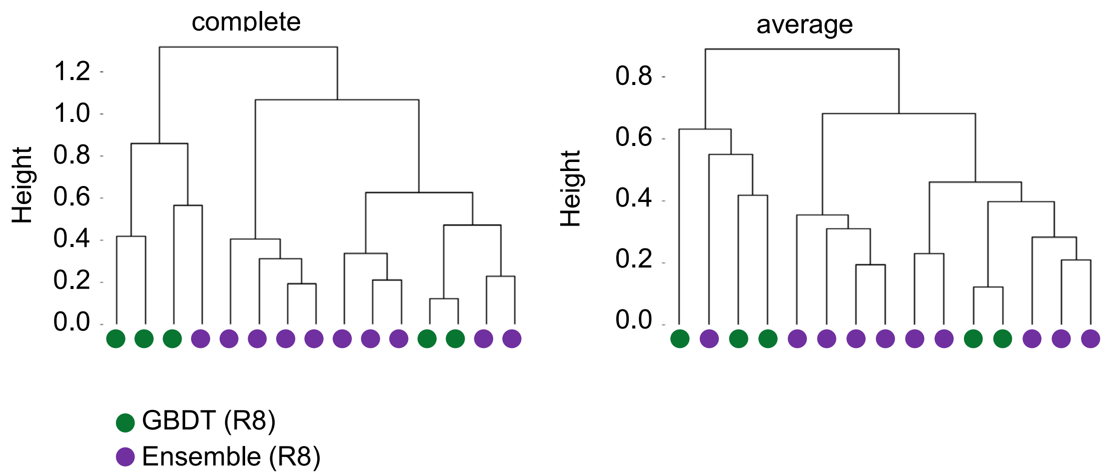


**Figure S4 Hierarchical clustering of the medium compositions predicted in R8**. The left and right panels show the complete and average methods used for clustering, respectively. Green and purple indicate the media predicted by the GBDT and Ensemble models, respectively.

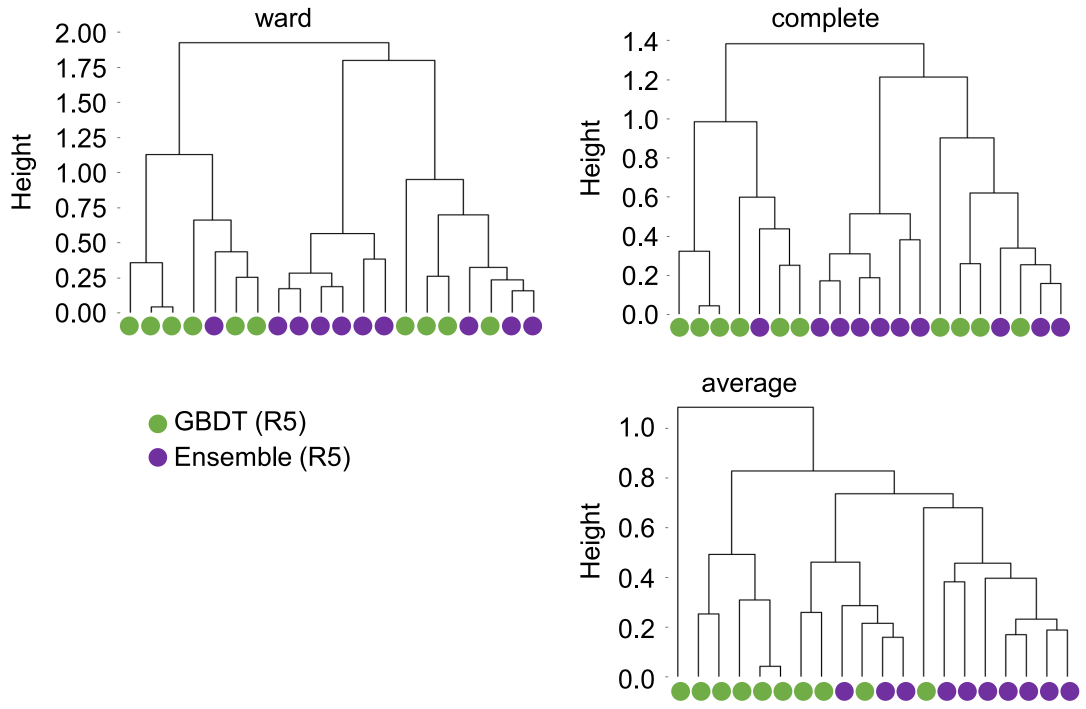


**Figure S5 Hierarchical clustering of the medium compositions predicted in R5**. The left, right, and bottom panels show the ward, complete, and average methods used for clustering, respectively. Green and purple indicate the media predicted by the GBDT and Ensemble models, respectively.

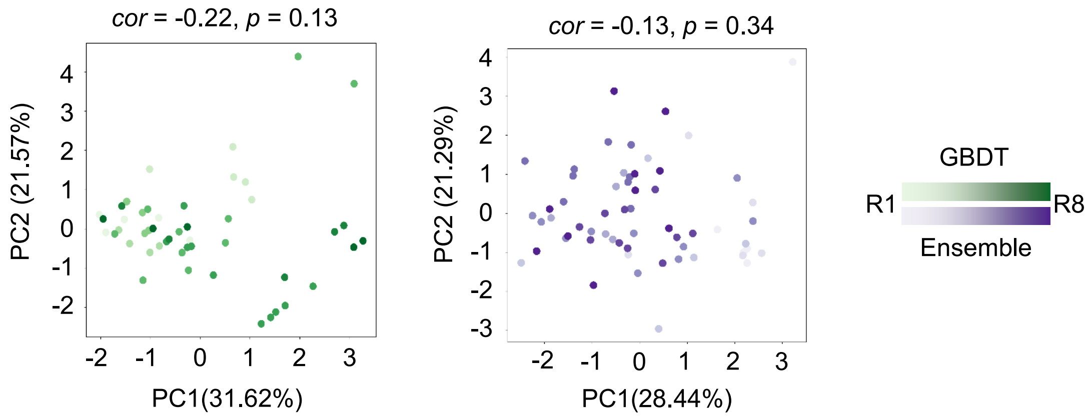


**Figure S6 PCA of the tested media from R1 to R8**. Green and purple indicate the media predicted by the GBDT and Ensemble models, respectively. Color gradation represents the varied rounds (R1~R8) of active learning. Spearman's correlation coefficients and p-values are indicated.

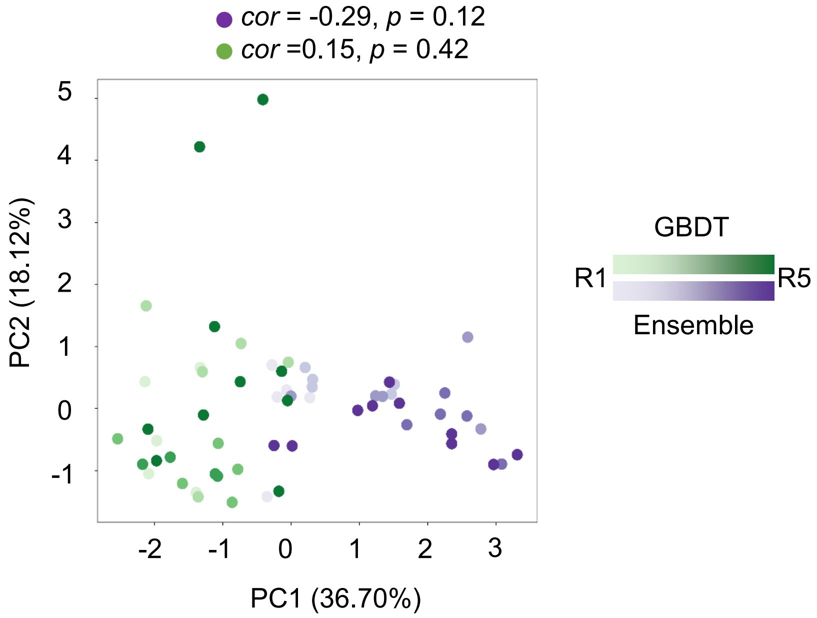


**Figure S7 PCA of the tested media from R1 to R5.** Green and purple indicate the media predicted by the GBDT and Ensemble models, respectively. Color gradation represents the varied rounds (R1~R5) of active learning. Spearman's correlation coefficients and p-values are indicated.

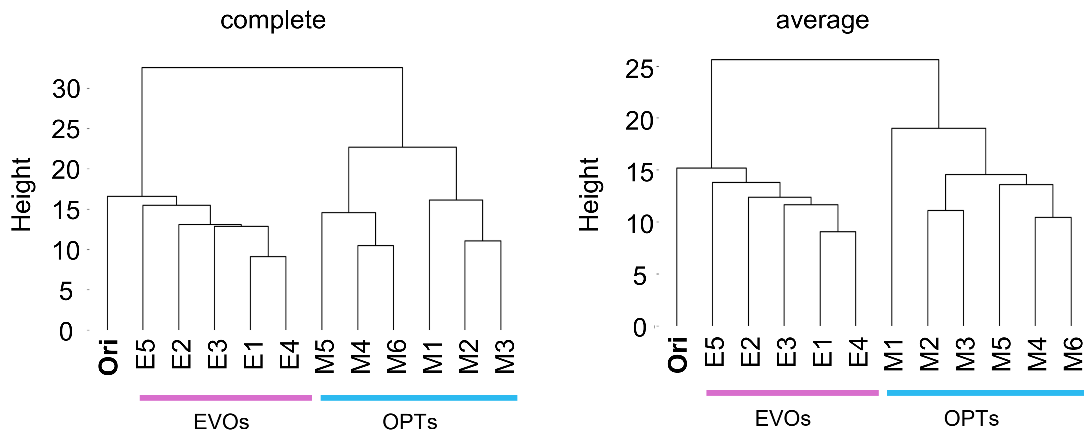


**Figure S8 Hierarchical clustering of transcriptomes.** The left and right panels indicate that the complete and average methods are used. Individual OPTs and EVOs are indicated.

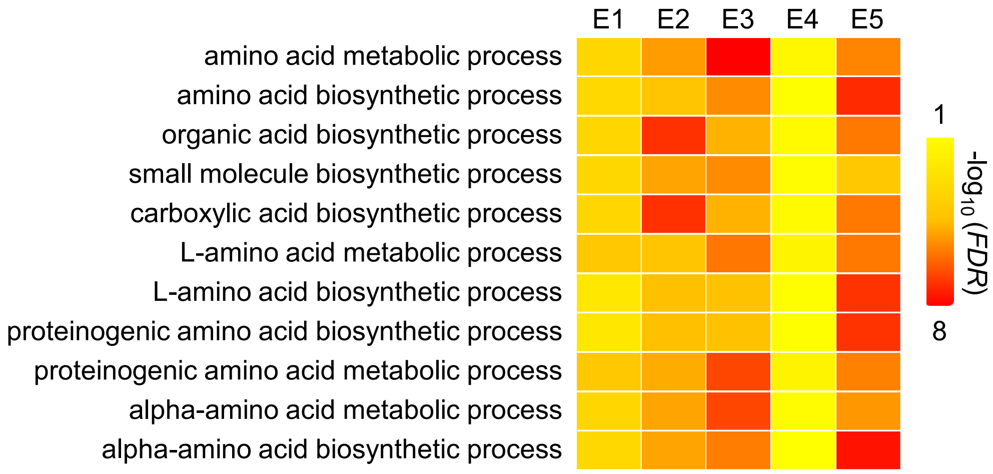


**Figure S9 Enriched GOs overlapped in all EVOs.** Color gradation represents FDR on a logarithmic scale.

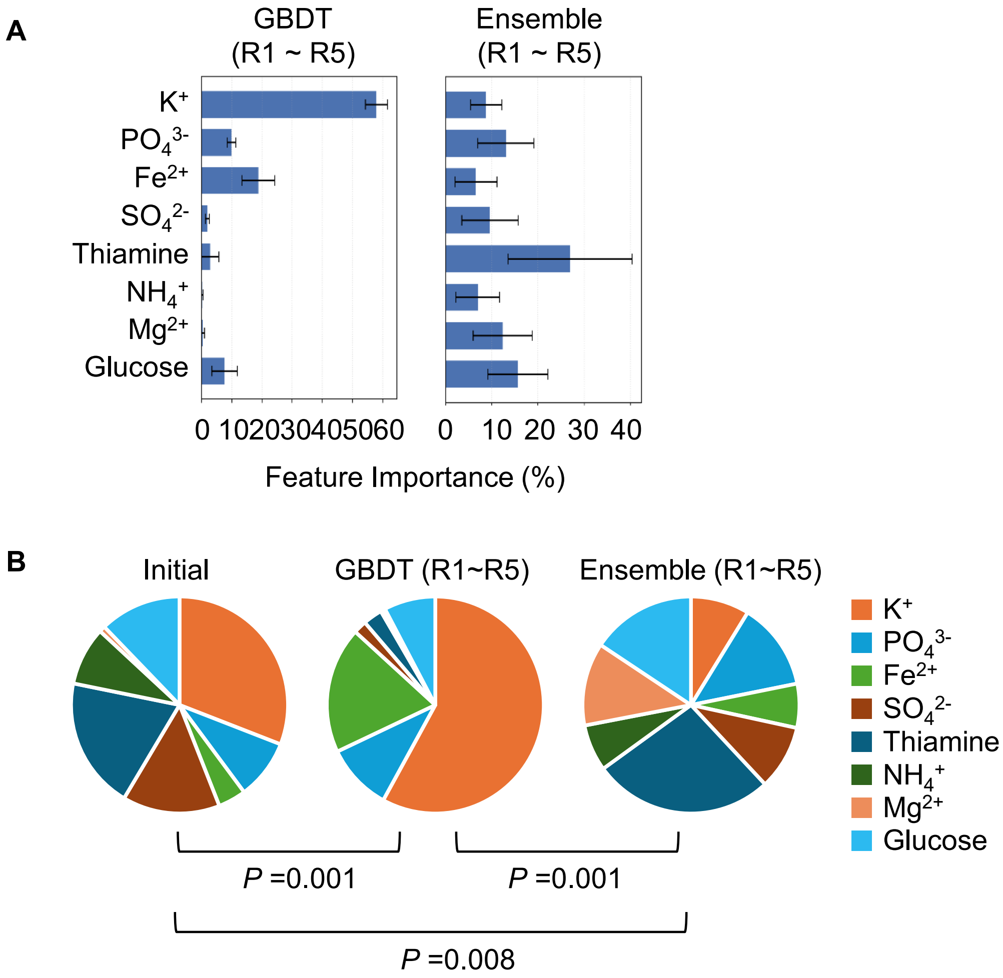


**Figure S10 Contributions of medium components to fitness increase. A.** Bar graphs of feature importance. Standard Errors indicate five repeated predictions. **B.** Pie graphs of feature importance. Initial, GBDT, and Ensemble indicate the predictions using the initial 99 media, the 30 GBDT-navigated media, and the 30 Ensemble-navigated media, respectively. Statistical significance of the pattern comparison is indicated (PERMANOVA).

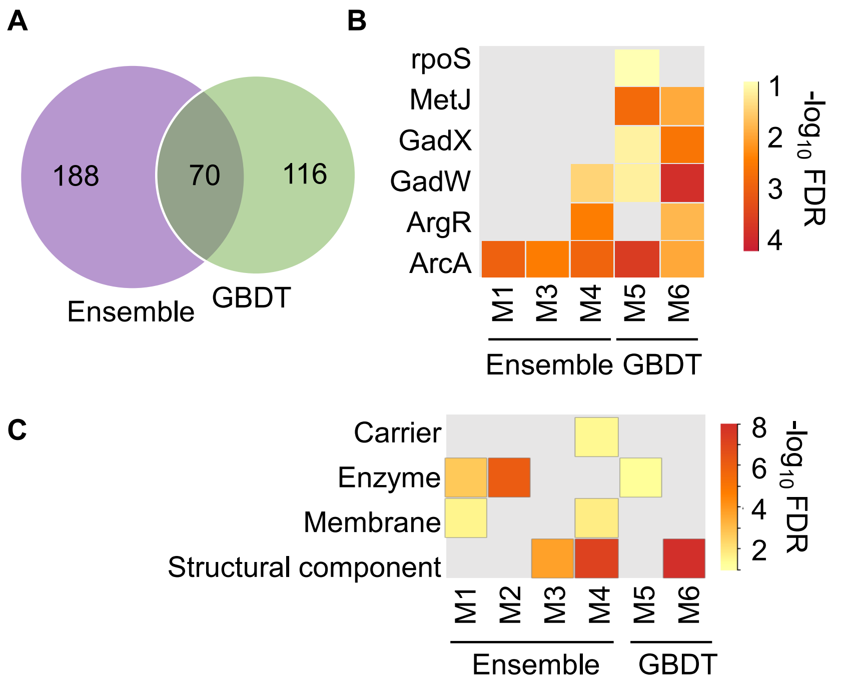


**Figure S11 DEGs and functional enrichment in OPTs. A.** DEGs identified in the Ensemble- and GBDT-derived OPTs. The numbers of overlapped and differentiated DEGs are indicated. **B.** Enriched regulons. **C.** Enriched gene categories. Color gradation represents FDR on a logarithmic scale.

**Table S1 Chemical compounds used for medium preparation.** Commercial product information, category, manufacturer, and concentration used to formulate the master mix are summarized.

| Pure chemical compounds | Commercial products | Categories |
|---|---|---|
| $KH_2PO_4$ | Dipotassium Hydrogenphosphate | inorganic salts |
| $K_2HPO_4$ | Potassium Dihydrogen Phosphate | inorganic salts |
| $FeSO_4$ | Iron(II) Sulfate Heptahydrate | inorganic salts |
| Thiamine (Vitamin B1) | Thiamin Hydrochloride | vitamins |
| $(NH_4)_2SO_4$ | Ammonium Sulfate | inorganic salts |
| $MgSO_4$ | Magnesium Sulfate Heptahydrate | inorganic salts |
| Glucose | D(+)-Glucose | sugars |
